# Resolution of recursive data corruption to transform T-cell epitope discovery

**DOI:** 10.64898/2026.03.30.710191

**Authors:** Grzegorz Preibisch, Michał Tyrolski, Piotr Kucharski, Stanislaw Giziński, Piotr Grzegorczyk, Sungho Moon, Sangwoo Kim, Balyn Zaro, Anna Gambin

## Abstract

Accurate prediction of MHC class I-presented peptides is essential for any vaccine or T-cell therapy design, yet reported gains on in silico benchmarks have not translated into clinical successes. Here we show that this discrepancy may come from a common methodological error: immunopeptidomics datasets are fundamentally contaminated by existing prediction models through prediction-based deconvolution and filtering, resulting in an iterative confirmation bias. An audit of the IEDB, the biggest database in the field, reveals that as of January 2025, 55.8% of assessable data are labeled by computational models rather than verified experimentally. This inflates in silico benchmarks while degrading real-world applicability on new data, effectively making it impossible to objectively test model performance, which can lead to choosing suboptimal solutions and decreasing the chance of any therapy’s clinical success. In silico simulation shows that iterative data corruption maintains high AUROC while top-of-list retrieval collapses. We reframe epitope discovery as a protein-centric learning-to-rank task and introduce deepMHCflare, a model evaluated exclusively on clean data. deepMHCflare achieves 0.80 Precision@4 on mono-allelic benchmarks versus 0.55-0.65 for gold-standard prediction models. A preclinical cancer vaccine study validated that 2 of the 4 deepMHCflare-nominated peptides were immunogenic, with a third independently confirmed in the literature.

---

Selecting peptide candidates for cancer vaccines and T-cell therapies remains a principal bottleneck in translational immunology [1–3]. Computational predictors are routinely used to rank peptides for synthesis and immunogenicity testing, yet improvements reported on public benchmarks have not consistently translated into higher prospective yields [2, 3]. This discrepancy is commonly attributed to biological complexity, but it also depends on how training and evaluation data are generated and curated.

Most immunopeptidomics experiments are performed on cells expressing multiple HLA alleles, and assigning each eluted peptide to its presenting allele requires computational deconvolution [4]. In practice, this deconvolution frequently relies on predictions from existing models, either explicitly, through prediction-guided pseudo-labelling [5], or indirectly, through iterative motif convergence [4]. As a result, peptide labels in public repositories become correlated with prior model outputs, as researchers were using those methods to label the data. Consequently, peptides matching existing expectations are preferentially retained, while discordant peptides are depleted. We term this recursive process *systematic confirmation bias* (Figure 1). Among current tools, the neural-network-based predictors NetMHCpan 4.1 [6, 7] and MHCflurry 2.0 [8, 9] exert influence through supervised deconvolution, while the motif-based MixMHCpred 3.0 [10, 11] operates through unsupervised motif convergence.

**Figure 1:**
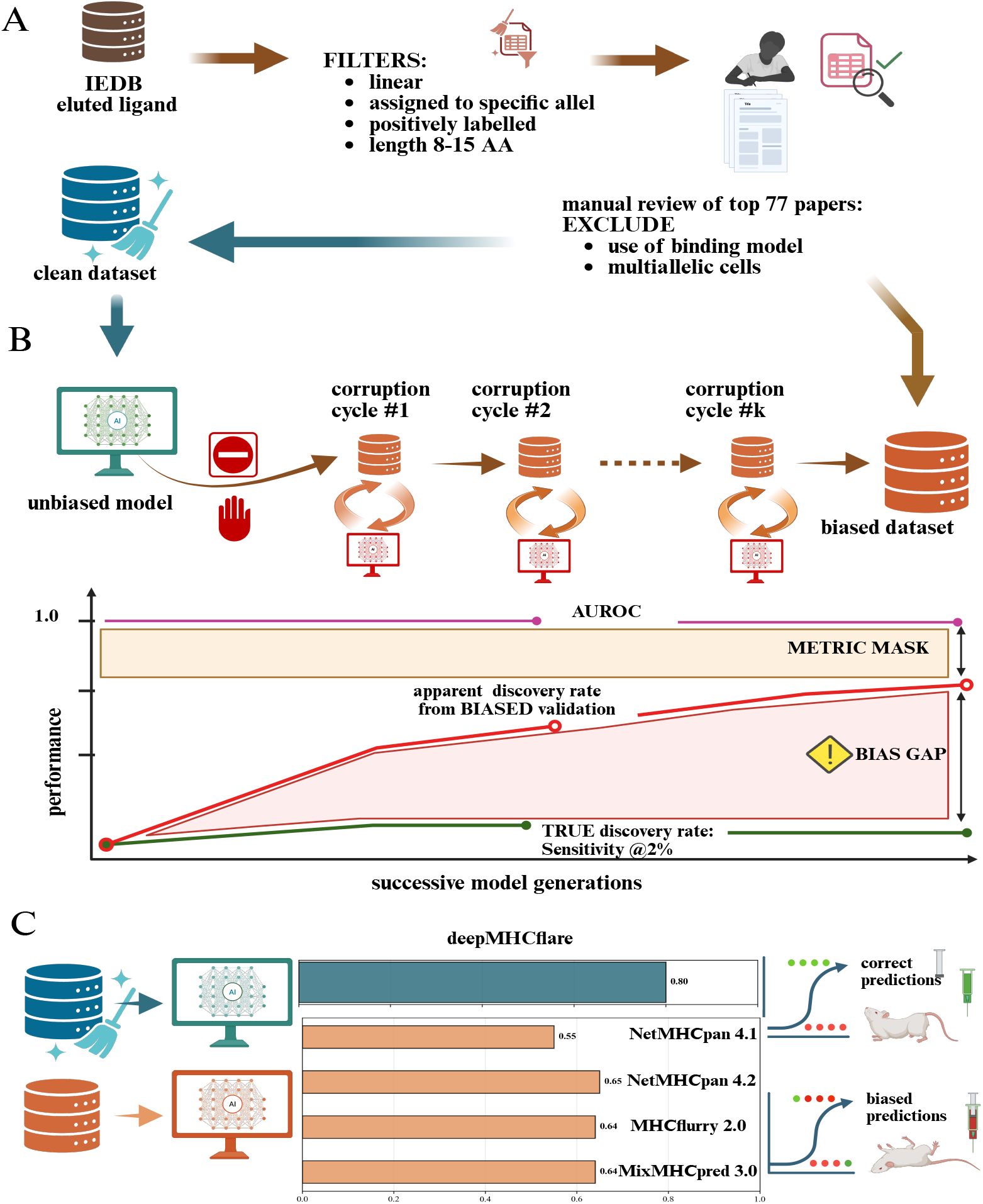
Systematic confirmation bias causes hidden performance stagnation. (A) The IEDB database was filtered according to the given criteria, and then, based on manual review of the publications that provided the largest amount of data, a clean dataset was constructed. The remaining part of the database is referred to as the biased dataset. (B) Consecutive cycles of model and data corruption and their performance. True Discovery Rate (green) stagnates at approximately 0.23 while Apparent Discovery Rate (red) rises to 0.58, creating a performance illusion (pink shaded area, called bisd gap). AUROC (magenta) remains above 0.89 throughout, masking the stagnation (yellow area). (C) The R-precision@4 achieved by DeepMHCflare and other (biased) models on on mono-allelic immunopeptidomics data. Created in https://BioRender.com

This problem is compounded by evaluation practice. Epitope discovery is a ranking problem under tight experimental budgets, where only a handful of top-ranked candidates per antigen can be tested [12]. Yet model assessment has relied predominantly on AUROC (area under the receiver operating characteristic curve), a global discrimination metric equivalent to the probability that a randomly chosen positive is scored higher than a randomly chosen negative. On a balanced dataset this is a natural measure of quality. However, in immunopeptidomics, where a single true binder must be found among 500 or more decoys, an AUROC of 0.95 means the true binder sits on average at position 25 – well outside the handful of candidates an experimentalist can afford to synthesise and test. AUROC is therefore insensitive to changes in top-of-list ordering on highly imbalanced datasets [13]. This effect is visible in the cancer vaccine case study (Figure 2C). Both models achieve similar AUROC, yet deepMHCflare concentrates true binders at the top of the ranking while NetMHCpan does not. Under predictor-dependent validation, AUROC can remain high even as enrichment among actionable candidates deteriorates (Figure 1B).

**Figure 2:**
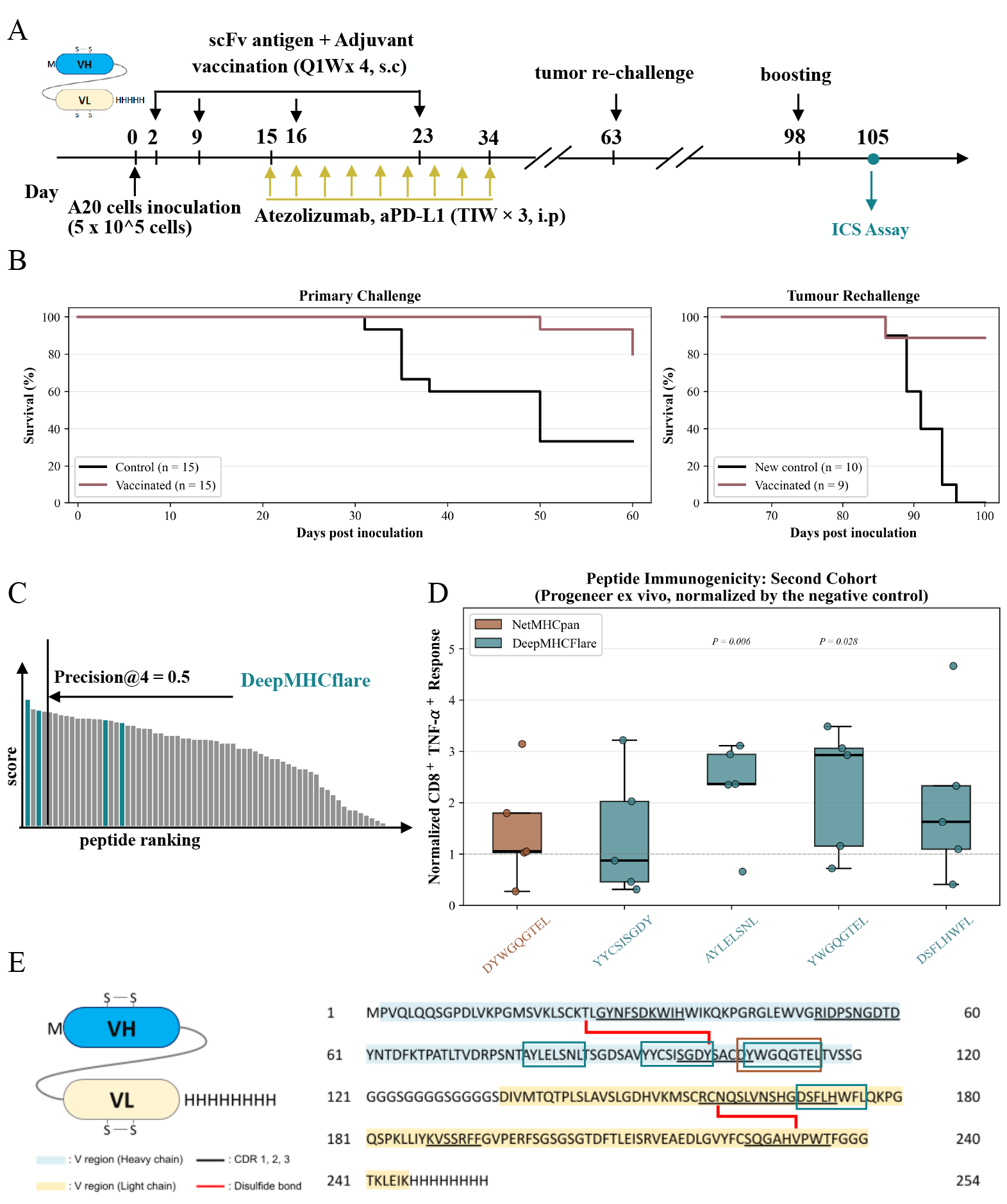
Preclinical cancer vaccine study. (A) Experimental timeline. BALB/c mice (*n* = 15 per arm) were inoculated with A20 cells, vaccinated with adjuvanted A20-scFv and Atezolizumab. Surviving mice were re-challenged on day 63, boosted on day 98, and splenocytes harvested for ICS on day 105 (*n* = 5). (B) Kaplan–Meier survival curves for primary challenge (*n* = 15 per arm) and tumour rechallenge (9 vaccinated survivors versus 10 naïve controls; log-rank *P* < 0.01 for both comparisons). (C) Illustrative peptide rankings for two models with similar AUROC; coloured bars mark validated epitopes, showing superior top-of-list enrichment by deepMHCflare. (D) Normalised CD8^+^ TNF-*α*^+^ responses (ICS, fold-change over negative control) for deepMHCflare-selected peptides. Exact *p*-values shown (one-sided paired *t*-test). (E) A20-scFv sequence with tested peptides boxed; heavy-chain (cyan) and light-chain (gold) variable domains and CDR loops shown in relevant colours.

Here, we present deepMHCflare, a protein-centric learning-to-rank predictor for MHC class I presentation, building on our earlier work [14]. We first audit the Immune Epitope Database (IEDB) [15] to quantify the prevalence of predictor dependence in public immunopeptidomics data. We then use in silico simulation to demonstrate how recursive data corruption decouples standard metrics from real predictive performance. Finally, we evaluate deepMHCflare exclusively on predictor-independent benchmarks and in a prospective cancer vaccine study, showing that the combination of predictor-independent training data, a protein-language-model backbone, and a ranking-aware training objective yields substantial improvements over established methods.

To quantify the extent of predictor dependence in public immunopeptidomics data, we audited all mass spectrometry-based eluted-ligand entries in the Immune Epitope Database (IEDB; January 2025 snapshot) [15], totalling 3,971,663 records. A two-stage protocol, programmatic SQL classification followed by systematic review of 37 publications, assigned each entry to one of four categories: *Clean* (experimentally resolved allele assignments from mono-allelic cell lines or allele-specific antibody pull-downs), *Biased* (allele labels assigned or confirmed by computational predictors), *Multi-allelic* (valid peptides from multi-allelic sources lacking allele-level resolution), and *Insufficient metadata* (entries whose provenance could not be determined). In total, only **44.2%** of assessable entries (1,511,463) were verified to be *clean* (Extended Data Tables 1a–c). Among the 3,417,839 entries where provenance could be determined, **55.8%** (1,906,376) carry predictor-dependent labels. The remaining *multi-allelic* and *insufficient*-metadata fractions, over one million entries each, cannot be used for allele-specific training without computational deconvolution, which reintroduces the same predictor dependence. These categories reflect fundamentally different degrees of predictor dependence in the provenance chain of each measurement. Based on this audit, we constructed a predictor-independent evaluation benchmark from the resolved mono-allelic subset [16, 17], enabling model selection and validation free from recursive contamination, cf. Figure 1A.

We designed an in silico simulation to quantify the effect of recursive data corruption on model performance (see Online Methods). Starting from a baseline model trained on clean mono-allelic data [16], we initiated an iterative cycle. Each generation’s model was used to filter an unlabeled dataset, and the filtered data, enriched with the model’s own biases, were used to train the next generation. This process was replicated five times with independent model initialisations to isolate the effect of data corruption from model training stochasticity, confirming that the observed performance degradation is driven by systematic bias rather than random variation in model training.

The results revealed a striking divergence as illustrated in Figure 1B. When evaluated on the corrupted data, models appeared to improve with each generation. AUROC remained above 0.89 across all iterations. To quantify the actual impact on discovery, we measured Sensitivity@Top2% – the fraction of the true peptidome that researchers can discover when applying the standard 2% rank threshold used by widely adopted prediction tools [6, 9] and commonly adopted by experimentalists selecting candidates for validation [18, 19] (see Online Methods). When evaluated against the clean, predictor-independent validation set, Sensitivity@Top2% declined substantially with each corruption iteration. Apparent discovery rate rose to 0.58 even as the true rate stagnated at 0.23, close to the 0.11 sensitivity expected from random selection on this validation set, cf. Figure 1B. High AUROC thus masked a near-complete collapse in the ability to retrieve actionable candidates. Together, biased data and a misleading metric create an illusion of progress. Models increasingly predict what was previously known rather than enabling new discoveries, and AUROC masks this stagnation [13].

We evaluated deepMHCflare against the most widely used prediction tools in the field, and the primary contributors to the data bias described above, NetMHCpan 4.1 [6], NetMHC-pan 4.2 [7], MHCflurry 2.0 [9], and MixMHCpred 3.0 [11], on a held-out set of mono-allelic cell lines [16]. Specifically, we measured performance using Precision@4, the fraction of true positives among the top four ranked candidates per protein. Such a measurement reflects the common experimental budget of selecting a small number of high-confidence candidates for validation. For deepMHCflare, training and evaluation used non-overlapping partitions of the same mono-allelic dataset, split by source antigen to prevent train–test leakage (see Online Methods). For competing models, such a guarantee cannot be made, as these datasets have been publicly available for several years and overlap with data sources used in their training pipelines. Therefore, any such overlap would favour the competing models, making our results a conservative estimate of the true performance difference. On this clean benchmark, deepMHCflare achieved 0.80 Precision@4, a 23-45% improvement over all established methods (Figure 1C).

We conducted additional ablation studies to assess generalisation. First, we held out 21 alleles entirely from training. The model retained meaningful ranking ability on these unseen alleles, confirming that it captures transferable binding patterns rather than memorising allele-specific features. Second, we tested generalisation on the multi-allelic HLA Ligand Atlas [20], an unseen, out-of-distribution dataset of over 90,000 ligands from 227 patient tissue samples. deepMHCflare generalised to this clinically realistic regime. We then retrained the model on additional clean data to quantify the improvement (Supplementary Note; see Online Methods).

To assess whether benchmark gains translate to a practical setting, we conducted a prospective cancer vaccine study in the A20 BALB/c murine lymphoma model (Figure 2A). Vaccinated mice showed significantly prolonged survival compared with controls in both the primary challenge and tumour rechallenge (Figure 2B). deepMHCflare ranked candidate peptides from an A20 B-cell lymphoma scFv antigen [21, 22] for the H-2K^d^ allele; despite comparable AUROC, deepMHCflare concentrated validated epitopes near the top of the ranked list (Figure 2C). The top four peptides were synthesized and tested for immunogenicity by intracellular cytokine staining (see Online Methods for details).

Two of four deepMHCflare-selected peptides elicited significant CD8^+^ TNF-*α*^+^ responses (*P* = 0.006 and *P* = 0.028, one-sided paired *t*-test; Figure 2D), and a third (YYCSISGDY) was independently reported as the only tumour-specific CDR3-derived epitope from A20 [21] (Figure 2E; Extended Data Table 6).

The confirmation bias we describe may extend beyond MHC prediction. Any setting in which model outputs shape subsequent training data is inherently susceptible to recursive contamination [23]. As computational methods become embedded in experimental pipelines across biology, systematic auditing of data provenance should become standard practice. Prospective validation across diverse antigens and HLA haplotypes will be required to establish the generality and practical impact of the effects observed here.

## Acknowledgements

We gratefully acknowledge Poland’s high-performance computing infrastructure PLGrid for providing computer facilities and support within computational grant no. PLG/2025/018310. We also acknowledge PCSS for providing access to the Eagle HPC cluster within computational grant no. pl0418-01. AG was supported by the National Science Centre, Poland, project 2024/53/B/ST6/03852.

## Author Contributions

G.P. conceived the learning-to-rank methodology, identified and quantified the recursive data bias, designed and executed the in silico bias-simulation experiments, trained production models, performed data analysis, participated in peptide selection for the prospective validation study, and led the writing of the manuscript. M.T. engineered the machine learning pipeline, developed the model architecture and training methodology, ran model training experiments, and contributed to writing and editing the manuscript. P.K. engineered the machine learning pipeline, provided technological oversight, and implemented evaluation metrics. S.Gi. engineered the machine learning pipeline, participated in peptide selection for the prospective validation study, contributed to training early model versions, and was involved in early methodology development. P.G. performed the final quantification of data bias in the IEDB audit. S.M. and S.K. designed and carried out the experimental vaccination and intracellular cytokine staining assays. B.Z. supervised experimental mass spectrometry methodology. A.G. supervised computational and statistical methodology.

## Competing Interests

G.P., M.T., P.K., P.G., and S.Gi. are or were employees of Deepflare. B.Z. and A.G. serve as scientific advisors to Deepflare. S.M. and S.K. are employees of Progeneer. Deepflare and Progeneer may benefit commercially from the methods described in this work.

## Data and Code Availability

Source code, trained model weights, and raw experimental data will be made available upon publication of the peer-reviewed article.

## Online Methods

### Data Curation and Quality Audit

The Immune Epitope Database (IEDB) [15] provides the foundational training data for virtually all MHC binding prediction algorithms and, equally importantly, serves as a primary source of evidence about specific epitopes. Any bias or error in the database therefore impacts both prediction tools and researchers who search the database directly, assuming it provides only experimental data. To quantify what fraction of the data may be impacted by bias, we developed a two-stage protocol - programmatic SQL classification followed by systematic paper-level review of 37 publications - that categorised entries into four groups:

- **Clean**. Data in which no prediction was used to assign an allele to a peptide. This group comprises immunopeptidomics data from mono-allelic cell lines and from studies that used allele-specific antibodies on multi-allelic samples (e.g. BB7.2, which precipitates only peptides bound to HLA-A2 [24]), and entries confirmed clean by paper-level review.
- **Biased**. Entries whose allele labels were assigned or confirmed by computational predictors, including multi-allelic data subjected to computational deconvolution, rank-threshold filtering, or prediction-based quality control.
- **Multi-allelic**. Valid peptides from multi-allelic sources labelled only at the class level (e.g., “HLA class I”), which require computational deconvolution for allele-specific use.
- **Insufficient metadata**. Entries whose allele-assignment method could not be determined from the available metadata - typically antibody-based captures where host genotype information is missing.

The systematic paper-level review of 37 publications reclassified 1,038,004 entries as biased and 823,151 as clean beyond what the programmatic SQL classification identified. Among the 3,417,839 assessable entries (Clean + Biased), 55.8% carry predictor-dependent labels (Extended Data Tables 1a,b). Classification verdicts, entry counts, and evidence quotes for each reviewed publication are provided in Extended Data Table 1c.

### Dataset description

We constructed a predictor-independent benchmark dataset free from the influence of existing model predictions through a multi-stage curation protocol applied to all eluted-ligand records classified as *Clean* in the IEDB as of January 2025. This dataset comprises 300,869 predictor-independent records from studies that passed all criteria, primarily from Sarkizova et al. [16] and Faridi et al. [17]. It was used both for validation of deepMHCflare and to simulate systematic confirmation bias.

In both cases, to avoid train–test leakage we separated data by holding out specific proteins from which peptides were or could have been derived. For deepMHCflare we additionally held out a set of alleles to perform ablation studies of the model’s generalisation to unseen alleles. Remaining records were retained as additional training data representative of publicly available, potentially contaminated data, as the clean subset was a small fraction of the total and the rigorous train–test split already held out a substantial number of alleles and peptides.

For multi-allelic validation, we selected patient tissue samples from the HLA Ligand Atlas [20], comprising over 90,000 ligands from 227 patient tissue samples across 51 alleles. These samples provided a realistic multi-allelic evaluation setting in which the model must simultaneously discriminate binders across all alleles expressed by a given individual.

### Evaluation metric

To simulate a budget-constrained setting where an experimentalist must select only four peptides, we define **Precision@4**. For each protein in the test set, we generated a candidate pool (1:512 positive-to-negative ratio) and ranked all candidates by predicted binding score on the mono-allelic benchmark. Precision@4 measures the fraction of true positives among the top four ranked candidates for each protein. For contexts with fewer than four positives, we use R-Precision (Precision@*R*, where *R* is the number of true positives) to avoid penalising the model for an impossible target.

### Learning-to-Rank Framework for Protein-Level Epitope Discovery

We reframed MHC class I presentation prediction as a protein-centric learning-to-rank task. For each experimentally validated epitope, we identified its source protein and generated a candidate pool of all possible 8–15-mer peptides using a sliding window. From this pool, 128–256 peptides served as negatives for each positive. This context-aware hard-negative sampling ensured that negatives included peptides highly similar to the positive-including single-residue extensions, truncations, and overlapping sequences-forcing the model to learn fine-grained features that distinguish true epitopes from near-identical decoys. All sampled negatives were cross-referenced against the full set of known positives to prevent false negatives.

The training objective combines a pairwise margin ranking loss, weighted by LambdaRank importance [25], with weighted binary cross-entropy. For each positive peptide *p* and negative peptide *n* from the same source protein, the margin ranking loss is:

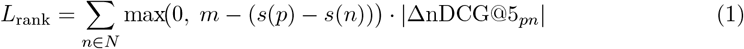

where *s*(·) are model scores, *m* is a fixed margin, and ΔnDCG@5_*pn*_ is the change in normalised discounted cumulative gain from swapping positions of items *p* and *n*, so that correctly ordering the top positions contributes most to the gradient. The combined loss is:

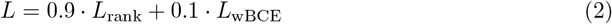

where *L*_wBCE_ is binary cross-entropy with class weights [0.1, 0.9], providing a stable classification signal under extreme class imbalance (∼0.2% positive rate).

The model was trained with the LAMB optimiser [26] at a static learning rate of 0.003 with weight decay of 0.2. The per-device batch size was 1024 with 64 gradient-accumulation steps, yielding an effective batch size of 65,536. Training used bf16-mixed precision with PyTorch DDP for 100 epochs, with validation every 5 epochs. Full hyperparameters are provided in Extended Data Table 3.

### Model architecture

deepMHCflare uses ESM2-t6-8M [27], a 6-layer, 8-million-parameter protein language model, as its backbone encoder. Each HLA allele is represented by a pseudo-sequence of approximately 182 amino acids comprising the alpha-1 and alpha-2 domains of the MHC class I heavy chain. Pseudo-sequences were extracted by pairwise alignment of full-length allele sequences against UniProt reference heavy chains using the biotite library [28]; the resulting database covers 14,856 HLA class I alleles. The MHC pseudo-sequence and candidate peptide (8–15 amino acids) are concatenated into a single input token sequence and processed jointly by the transformer encoder, so that the self-attention mechanism operates over both MHC and peptide positions.

Three pooling strategies are applied to the encoder’s final-layer hidden states: mean pooling with square-root length normalisation, element-wise max pooling, and the [CLS] token embedding. The resulting vectors are concatenated into a 960-dimensional representation and passed through a linear classification head (960 → 2) with sigmoid activation. Hidden dropout is set to 0.2; attention dropout is set to 0.0. The architecture is depicted in Extended Data Figure 3.

### Simulation of systematic confirmation bias

The predictor-independent dataset was partitioned into *K*=4 non-overlapping folds based on MHC allele (23 alleles per fold), with no allele overlap between folds. Within each fold, data were split into training (80%) and validation (20%) subsets by holding out separate protein sources to prevent peptide-level leakage. Negative samples were sampled from the same source antigens as the positives at a fixed positive-to-negative ratio of 1:4 for training and 1:9 for validation, approximating the class balance used by established predictors. The procedure was replicated five times with independent model initialisations to isolate the effect of data corruption from model training stochasticity, confirming that the observed performance degradation is driven by systematic bias rather than random variation in model training.

The simulation proceeded through four iterations (*i* ∈ {0, 1, 2, 3}). Iteration 0 served as the clean baseline: model *M*_0_ was trained on uncorrupted Fold 0. In each subsequent iteration: (1) model *M*_*i*_ −_1_ predicted binding scores for Fold *i*; (2) a positive observation retained its label only if its predicted rank fell within the top 2% of all scores for that fold, and all others were relabelled as negative-this cutoff mirrored the standard rank threshold employed by NetMHCpan 4.1 [6] and reflected filtering criteria documented in several large IEDB datasets; (3) the corrupted training data from Fold *i* was merged with the complete training set from iteration *i*−1:

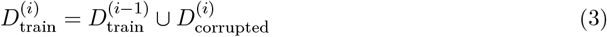

(4) a new model *M*_*i*_ was trained on the accumulated dataset. Each model was evaluated on both the original predictor-independent validation set and a corrupted validation set created by applying the same corruption rule. To quantify the impact of the bias caused by the 2% threshold present in the vast majority of biased papers, we evaluated what fraction of the peptidome may be discovered by the model at that threshold. We define this metric as **Sensitivity@Top2%**: the fraction of true positives whose predicted rank falls within the top 2% of all candidates.

### Prospective Validation Study Design

The prospective cancer vaccine study was conducted by Progeneer Inc. (Seoul, Republic of Korea). Candidate 8–9-mer peptides were predicted from the A20 B-cell lymphoma scFv antigen sequence using deepMHCflare. The top four ranked peptides were selected for testing. DYWGQGTEL, the top-ranked peptide predicted by NetMHCpan 4.1 [6], was additionally included as a reference control; notably, Warncke et al. [21] showed that this peptide induces suppressive CD4^+^ rather than cytotoxic CD8^+^ T-cell responses. Eight-week-old female BALB/c mice were inoculated subcutaneously with 5 × 10^5^ A20 cells on day 0. The vaccinated group (*n* = 20) received four weekly subcutaneous doses of A20-scFv protein (50 µg) formulated with ProLNG-002 adjuvant (TLR4a 5 µg + TLR7/8a 70 µg; days 2, 9, 16, 23) combined with Atezolizumab administered intraperitoneally three times per week for three weeks (days 15–34). Vehicle controls received no treatment.

By day 34, tumor volumes in the control and vaccinated groups showed a statistically significant difference (Mann–Whitney test, p < 0.0001), demonstrating the vaccine’s antitumor efficacy. At day 63, surviving mice were re-challenged with A20 cells in the contralateral flank; all re-challenged mice remained tumour-free, with an additional control group showing no survival. This confirmed a durable antitumour memory response. To identify which peptides drive this T-cell response, surviving mice received a single vaccine boost on day 98. One week later (day 105), splenocytes were harvested and stimulated with the four deepMHCflare-selected peptides and DYWGQGTEL as a reference control.

Spleens were dissociated using the Spleen Dissociation Kit (Miltenyi Biotec, 130-095-926) following the manufacturer’s protocol. Tissues stored in RPMI-1640 were transferred to C-tubes and mechanically dissociated into single-cell suspensions using the GentleMACS Dissociator (Miltenyi Biotec). Cell suspensions were filtered to remove debris and treated with RBC lysis buffer at room temperature for 30 minutes. After washing, cells were resuspended in RPMI-1640 supplemented with 10% FBS, counted using a 3-FL counter (ThermoFisher).

Single-cell suspensions (1 × 10^6^ cells/100 µL) were seeded into 96-well plates and stimulated with specific peptides (1 µg/mL each; Genscript) in the presence of Monensin (1:1000; Invitrogen, 00-4505-51) and Brefeldin A (1:1000; BioLegend, 420601) in RPMI-1640 medium (Welgene, LM011-03) supplemented with 25 mM HEPES, 50 µM 2-mercaptoethanol, and 10% fetal bovine serum (FBS; Gibco, 16000-044). Cells were incubated for 6 hours at 37°C in a 5% CO_2_ incubator. After stimulation, cells were stained with surface markers CD3 (FITC; ThermoFisher, 11-0032-82), CD4 (PE-Cy7; ThermoFisher, 25-0042-82), and CD8 (PerCP-Cy5.5; ThermoFisher, 45-0081-82) in Flow Cytometry Staining Buffer (Invitrogen, 00-4222-26) for 30 minutes at 4°C in the dark. Cells were then washed twice and fixed/permeabilized using the Intracellular Fixation & Permeabilization Buffer (Invitrogen, 88-8824-00) for 30 minutes. Intracellular cytokine staining was performed with antibodies against IFN-*γ* (PE; ThermoFisher, 12-5773-82), IL-2 (APC; ThermoFisher, 17-7021-82), and TNF-(APC; ThermoFisher, 17-7321-82) for 20 minutes at room temperature in the dark. After washing, cells were resuspended in permeabilization buffer and analyzed using a flow cytometer (Agilent, Novocyte3000).

### Immunogenicity analysis

Immunogenicity was assessed per peptide by one-sided paired *t*-test comparing the percentage of TNF-*α*-producing CD8^+^ T cells in peptide-stimulated versus matched negative-control (DMSO) samples from the same mouse (*p* < 0.05). This was the single pre-specified primary readout. No correction for multiple comparisons was applied; uncorrected *p*-values are reported for each peptide independently, consistent with standard practice in small-cohort epitope immunogenicity studies.

Immunogenicity was read out on day 105, after tumour re-challenge at day 63 and vaccine boost at day 98 (*n* = 5 vaccinated tumour-free survivors). Because these mice were selected as tumour-free survivors, they may represent strong immune responders; however, the within-mouse paired design controls for this enrichment. Four deepMHCflare-selected peptides and one reference peptide (DYWGQGTEL) were tested. Shapiro–Wilk tests confirmed that paired differences were consistent with normality for all peptides and controls (DMSO control: *p* = 0.52; formic acid control: *p* = 0.46), supporting the use of the paired *t*-test:

- **AYLELSNL** - 2.3×, *p* = 0.006. Significant.
- **YWGQGTEL** - 2.3×, *p* = 0.028. Significant.
- **YYCSISGDY** - 1.4×, not significant. Warncke et al. [21] independently identified YYCSISGDY as the only genuinely tumour-specific CDR3 peptide from A20 with no normal immunoglobulin homology, and reported that it induced antigen-specific effector T cells. Despite this literature support, the peptide did not reach significance in our assay.
- **DSFLHWFL** - inadvertently reconstituted in formic acid instead of DMSO. Against the matched formic-acid negative control: 2.0×, *p* = 0.170 (not significant). Against the DMSO negative control used for the other peptides: *p* = 0.053 (borderline, but this cross-vehicle comparison was not pre-specified). We conservatively classify DSFLHWFL as a weak positive given the confounded vehicle.

Secondary readouts (CD4^+^/CD8^+^ × IFN-*γ*, IL-2, Granzyme B) are reported in Extended Data Table 6. For bias-simulation experiments, standard errors were computed across five independent replications.

## Extended Data

**Extended Data Table 1a:** Headline classification of IEDB eluted-ligand entries (assessable subset).

| Category | Count | % of Assessable |
| --- | --- | --- |
| **CLEAN** | 1,511,463 | 44.2% |
| **BIASED** | 1,906,376 | 55.8% |
| **Assessable total** | **3,417,839** | **100.0%** |
| Multi-allelic unassigned | 1,351,486 | - |
| Insufficient metadata | 1,078,637 | - |
| **Grand total** | **3,971,663** |  |

**Extended Data Table 1b:**
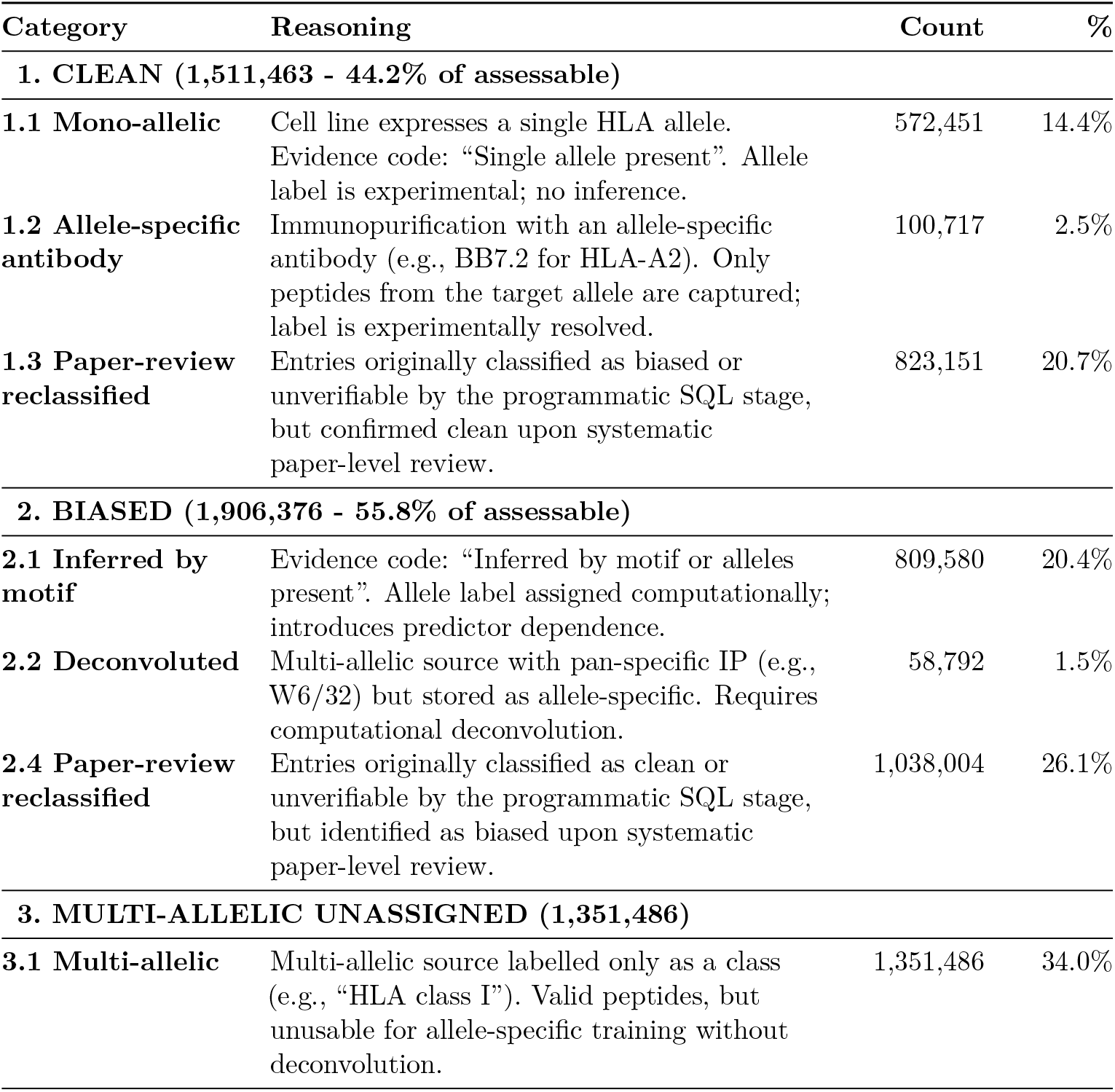
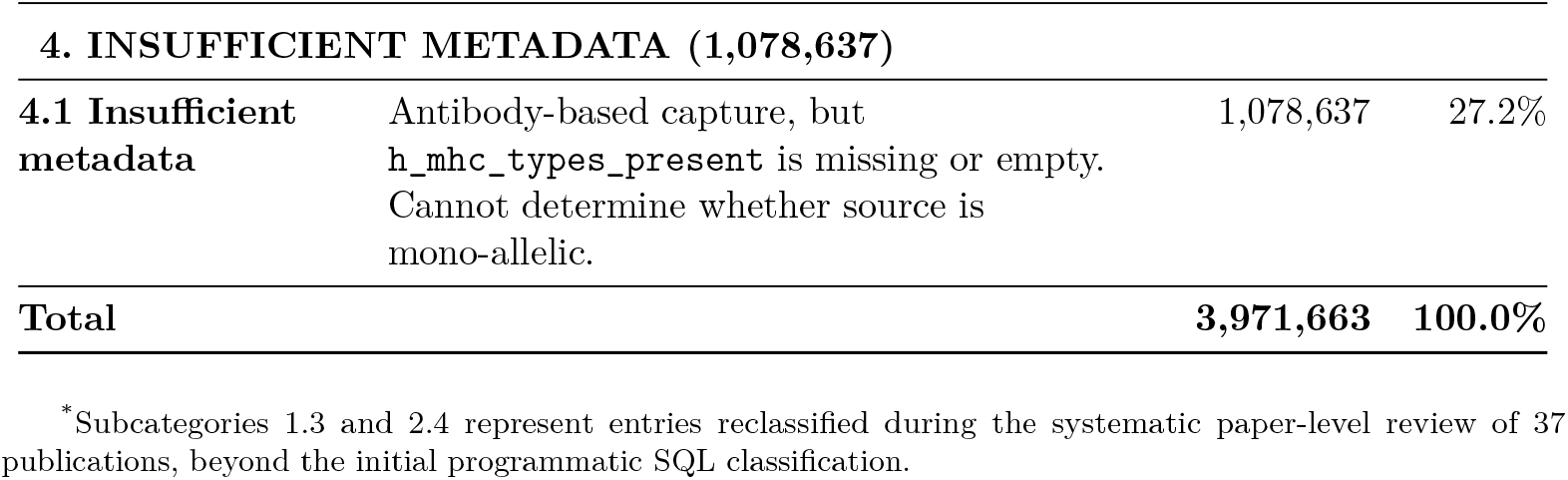
Detailed subcategory definitions and counts for each data usability classification.

**Extended Data Table 1d:** Specificity and IEDB counts for common immunopurification antibodies.

| Antibody | Specificity | Count |
| --- | --- | --- |
| **Pan-HLA Class I (Requires Deconvolution)** |  |  |
| W6/32 | Pan-HLA-I (A, B, C) | 844,216 |
| **Pan-HLA Class II (Requires Deconvolution)** |  |  |
| Tu39 | Pan-HLA-II (DR, DP, DQ) | 447,539 |
| B7/21 | HLA-DP | 86,438 |
| LB3.1 | HLA-DR | 71,172 |
| L243 | HLA-DR | 59,897 |
| **Allele-Specific (Clean Data)** |  |  |
| BB7.2 | HLA-A2 (A*02 family) | 52,688 |

**Extended Data Table 1c:**
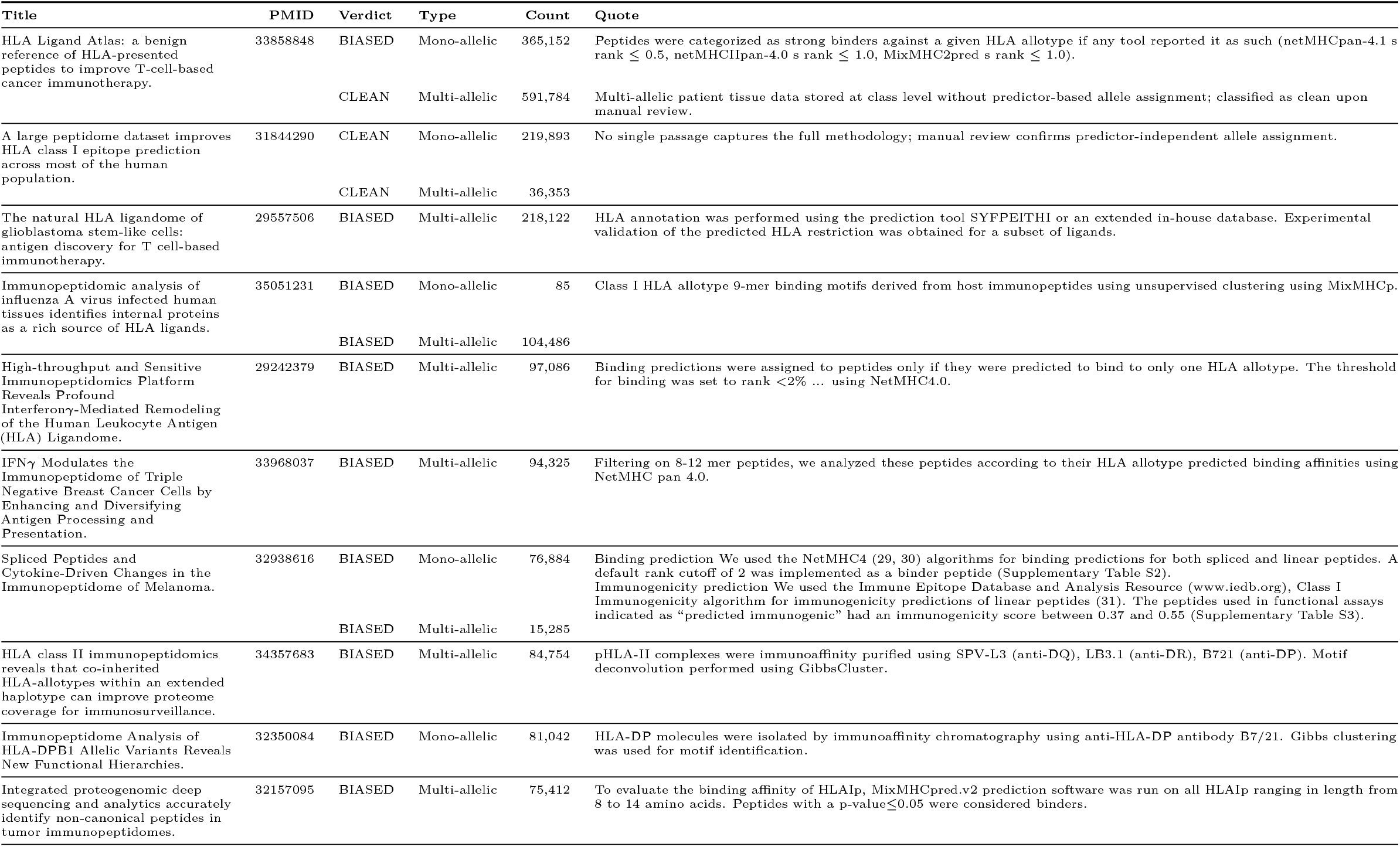
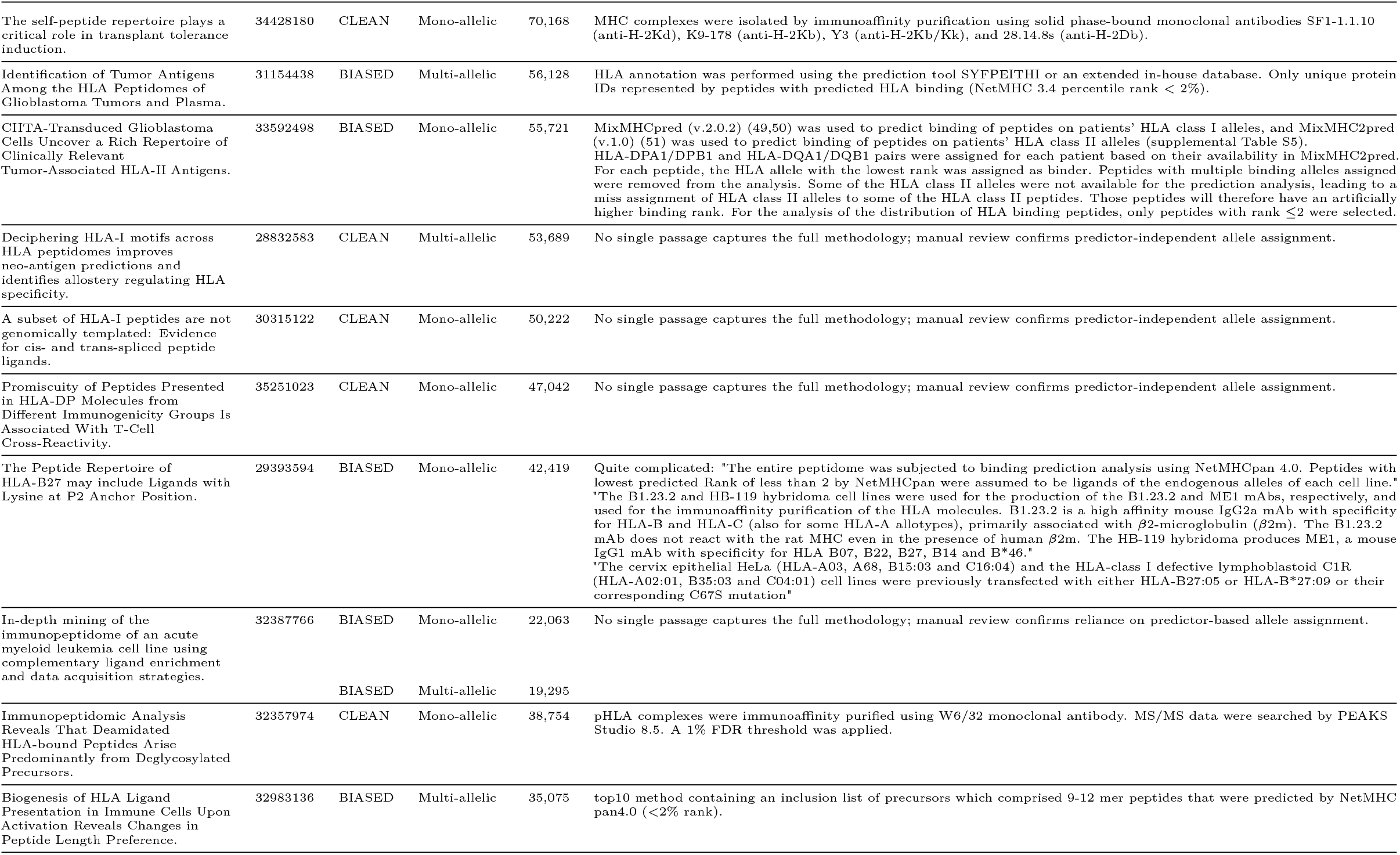
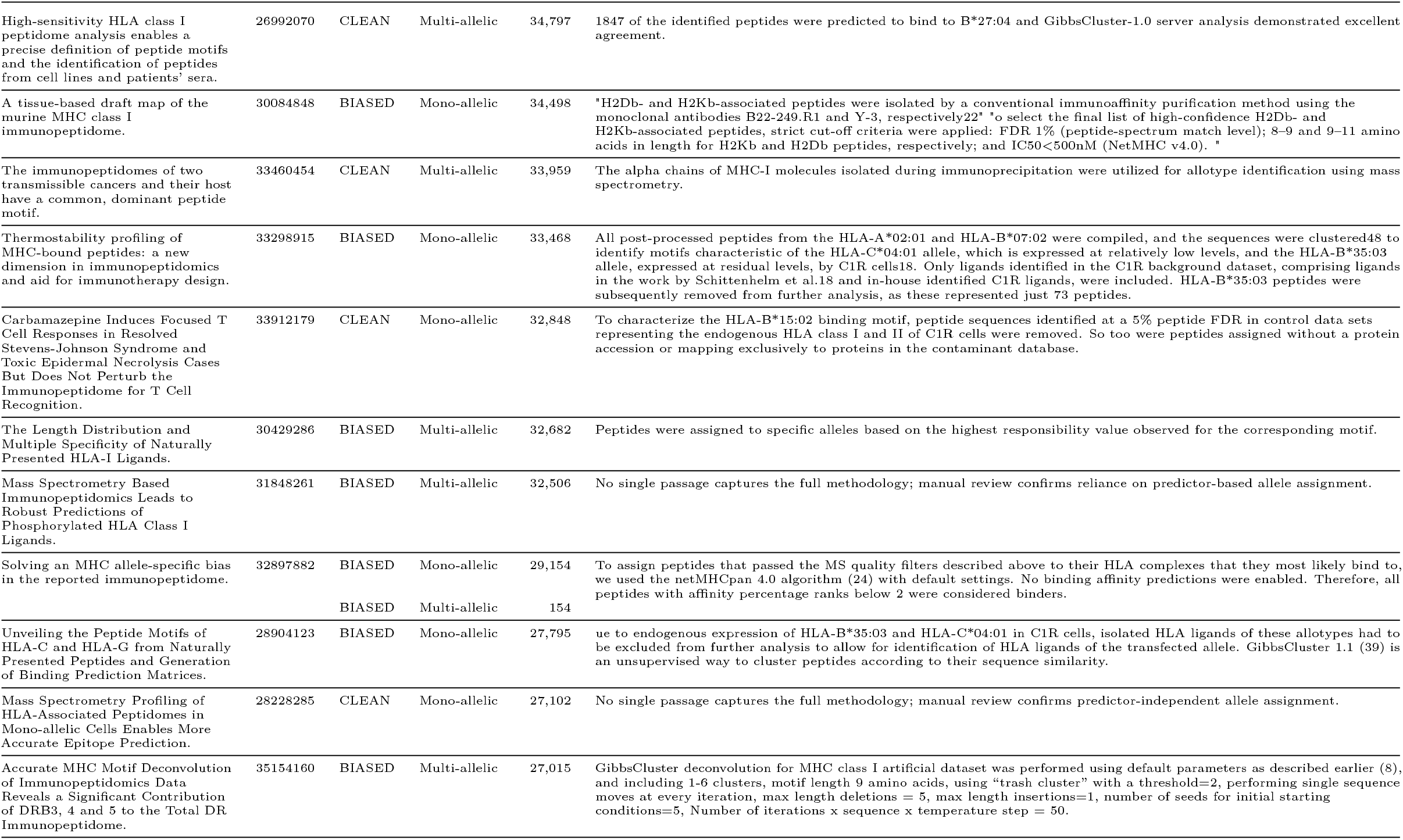
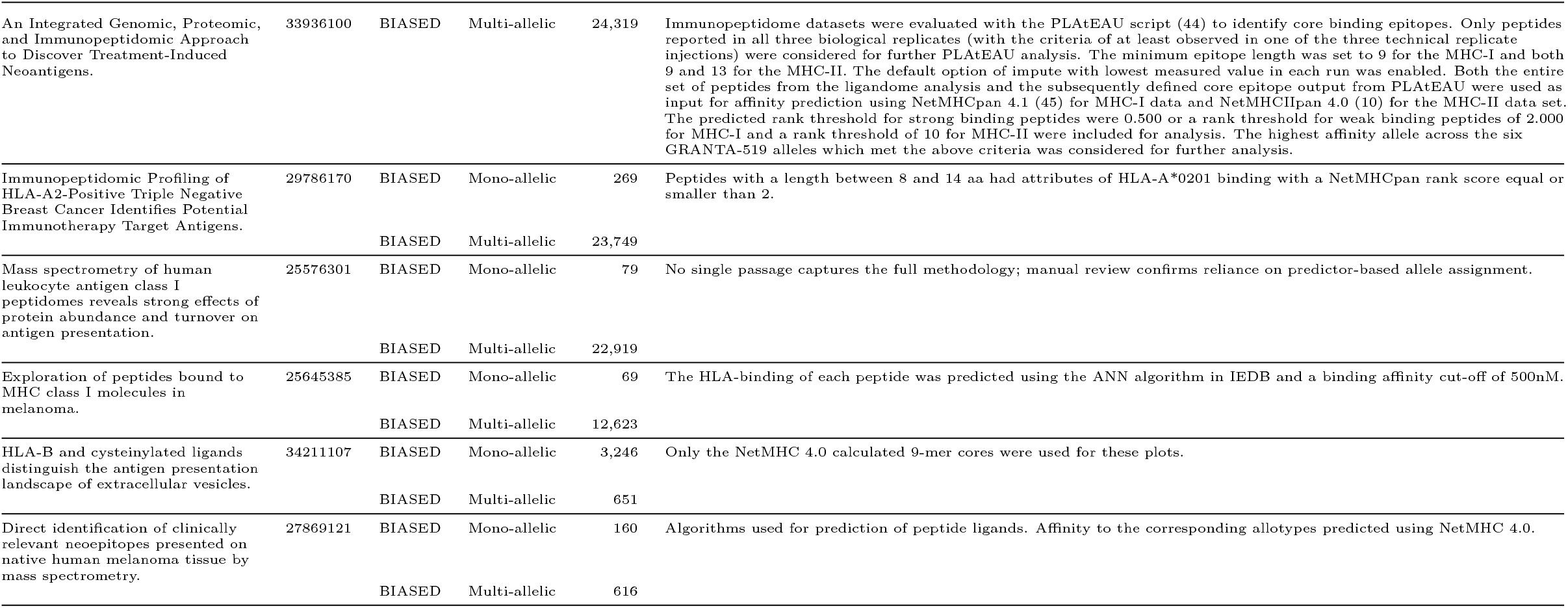
Systematic review of 37 publications with classification verdicts, entry counts, and evidence quotes.

**Extended Data Table 2:** Allele-to-fold assignment for the in silico bias simulation. The partition is fixed across all five replications (*n* = 3 … 7). Each fold contains 23 alleles.

| Fold | Alleles |
| --- | --- |
| 0 | A*02:04, A*30:01, A*30:02, A*33:01, A*33:03, B*08:01, B*18:01, B*35:01, B*38:01, B*40:06, B*42:01, B*45:01, B*49:01, B*50:01, B*52:01, B*58:02, C*02:02, C*03:03, C*04:01, C*04:03, C*05:01, C*07:04, C*08:01 |
| 1 | A*02:02, A*02:03, A*02:07, A*02:11, A*11:01, A*25:01, A*29:02, A*34:02, A*36:01, A*66:01, A*68:02, B*13:01, B*15:01, B*15:02, B*15:03, B*27:05, B*38:02, B*40:01, B*46:01, B*53:01, B*58:01, C*14:02, G*01:04 |
| 2 | A*02:01, A*02:05, A*03:01, A*11:02, A*23:01, A*24:02, A*24:07, A*26:01, B*07:02, B*13:02, B*15:10, B*15:17, B*40:02, B*44:02, B*44:03, B*51:01, B*55:02, B*56:01, B*57:03, C*03:02, C*07:02, C*08:02, C*17:01 |
| 3 | A*01:01, A*02:06, A*31:01, A*32:01, A*34:01, A*68:01, A*74:01, B*07:04, B*14:02, B*35:03, B*35:07, B*37:01, B*54:01, B*55:01, B*57:01, C*01:02, C*12:02, C*12:03, C*14:03, C*15:02, C*16:01, G*01:01, G*01:03 |

**Extended Data Table 3:**
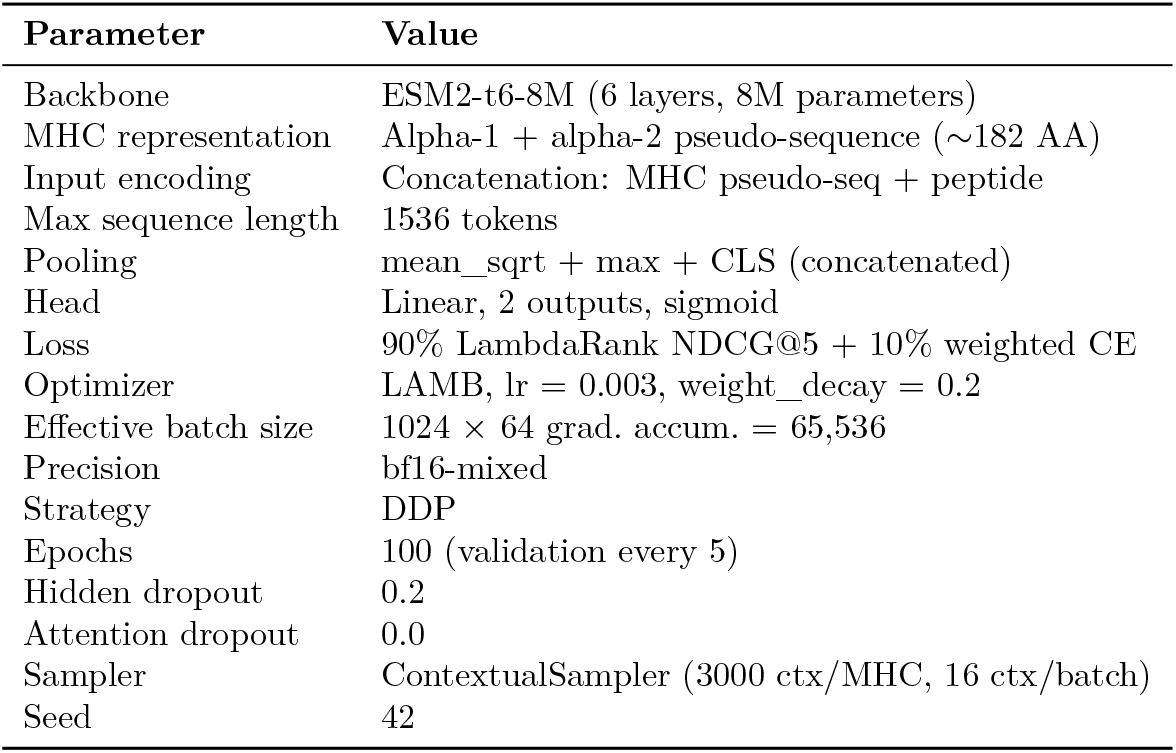
Hyperparameters for the final deepMHCflare model.

**Extended Data Figure 1:**
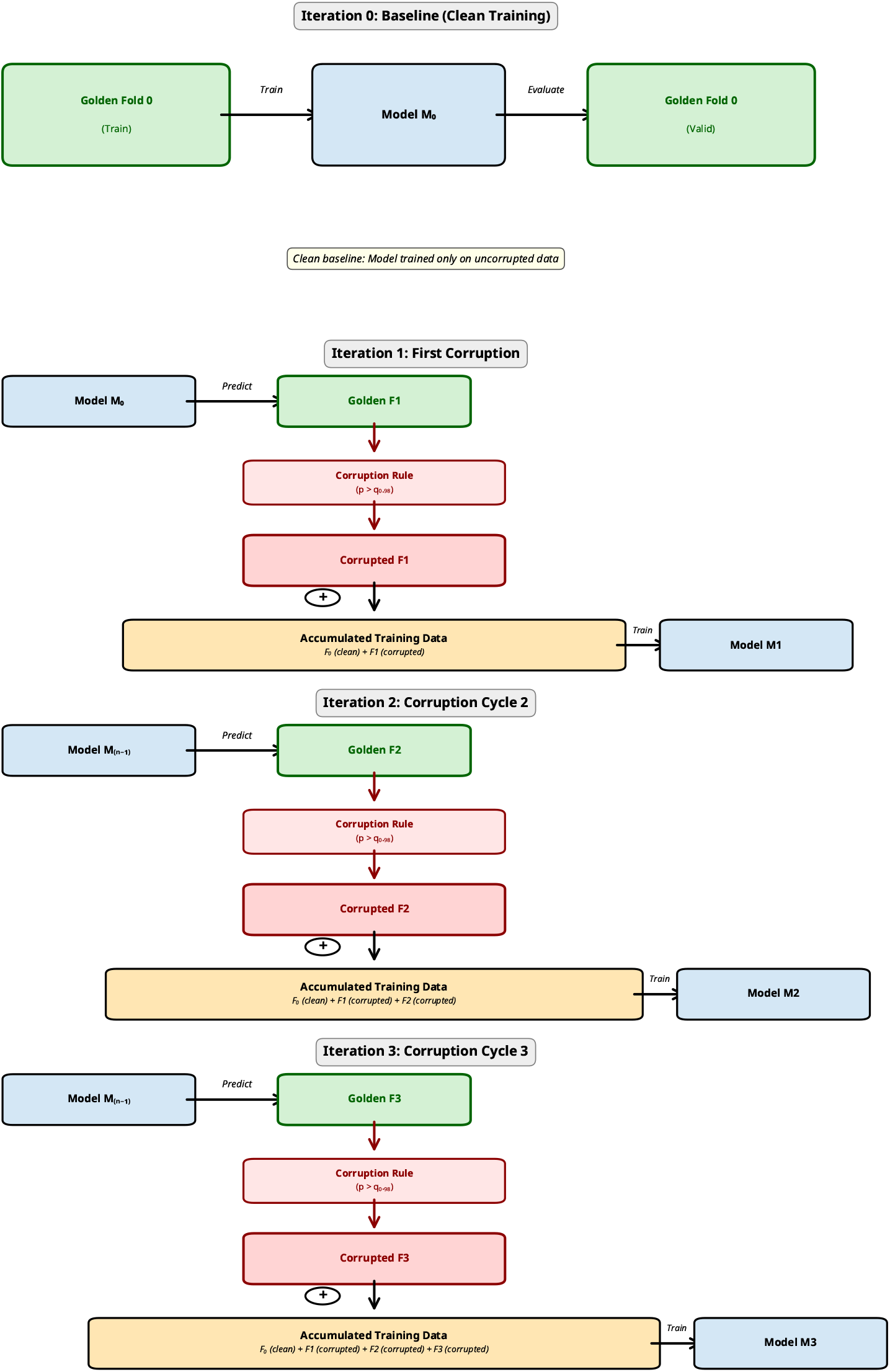
Iterative simulation of systematic confirmation bias. The four-iteration process used to simulate bias accumulation. In each iteration *i* > 0, the model from the previous step (*M*_*i*_ − _1_) predicts on and subsequently corrupts the next clean fold, which is then added to the growing training dataset.

**Extended Data Figure 2:**
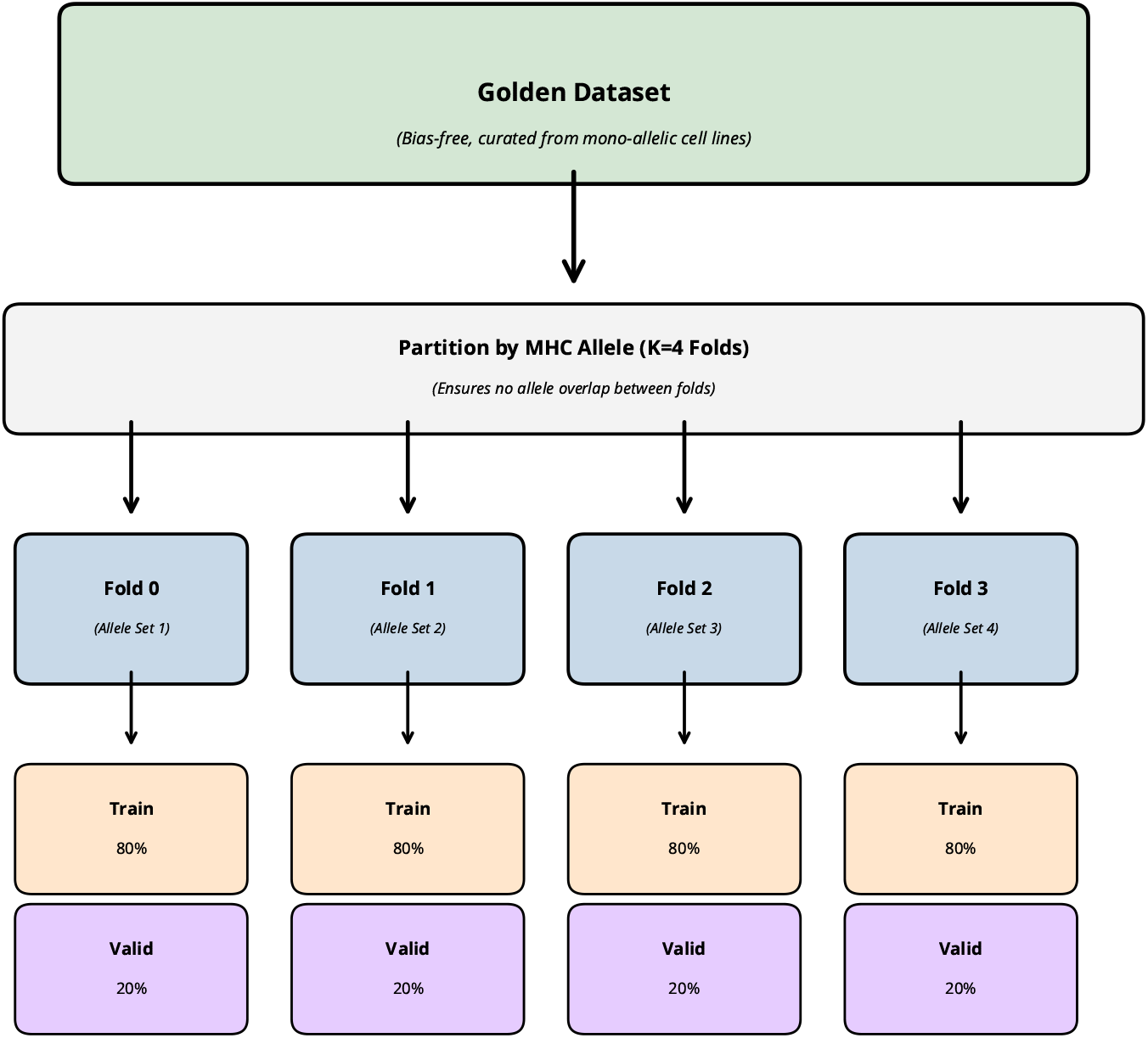
Predictor-independent dataset partitioning strategy. The predictor-independent dataset was partitioned into four non-overlapping folds defined by MHC allele. Within each fold, data were further split into 80% training and 20% validation sets.

**Extended Data Table 4:** Design dimensions explored during model development.

| Dimension | Range explored | Final choice |
| --- | --- | --- |
| Architecture | Single encoder vs. dual (separate MHC/peptide) encoders, incl. funnel variant | Single encoder (concat) |
| Backbone | ESM2 variants (8M–650M) | ESM2-t6-8M |
| Loss function | Pure CE, LambdaRank (Precision@$k$, NDCG@$k$ for $k=5,10$), combinations at ratios 0.1–0.9 | 90% LambdaRank NDCG@5 + 10% weighted CE |
| Learning rate | $10^{-6}$ to $10^{-3}$ | $3 \times 10^{-3}$ |
| Batch size | 64–1024, grad. accum. 1–128 | $1024 \times 64 = 65\text{K}$ effective |
| Regularisation | Hidden dropout 0.0–0.2, weight decay 0.05–0.3, layer freezing 0–5 layers | dropout = 0.2, weight_decay = 0.2, no freezing |
| Sampling | Contextual (50–3000 ctx/MHC), parent-protein, mixed-batch | Contextual, 3000 ctx/MHC, 16 per batch |
| Scheduler | None, CosineAnnealing, custom TwoLRs | None (static LR) |
| Datasets | 7+ dataset generations, interspecies, golden subsets | Predictor-independent mono-allelic + curated multi-allelic |

**Extended Data Figure 3:**
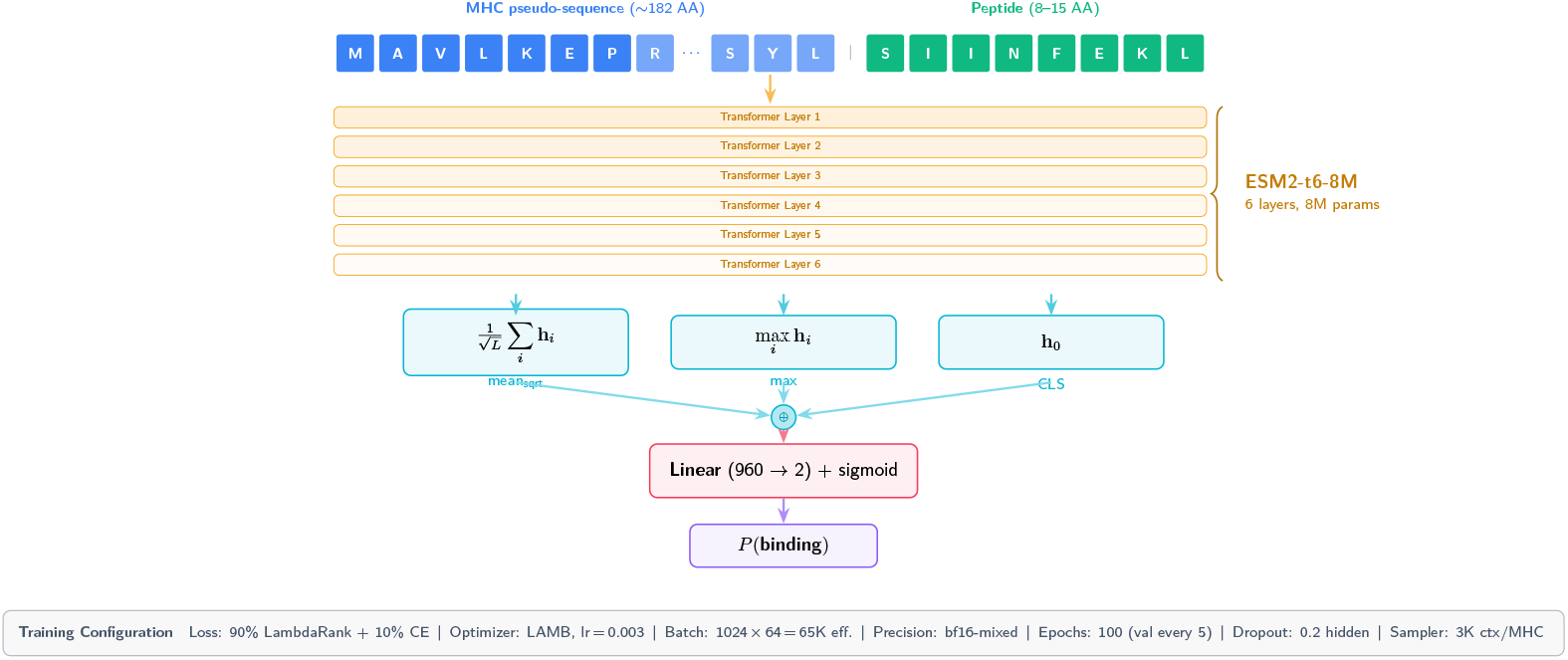
deepMHCflare architecture. The MHC alpha-1/alpha-2 pseudo-sequence (∼ 182 AA) is concatenated with the candidate peptide (8–15 AA) and processed by a pretrained ESM2-t6-8M encoder (6 transformer layers, 8M parameters). Three pooling strategies - mean with square-root normalisation, max, and CLS token - produce a 960-dimensional representation passed through a linear head to yield a binding probability.

**Extended Data Table 5:**
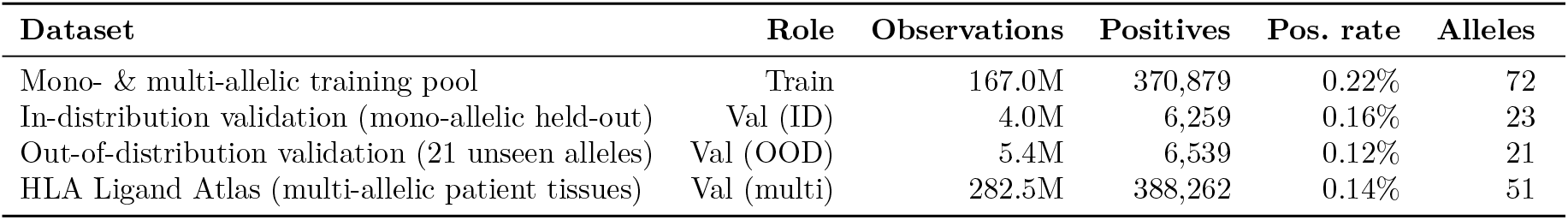
Summary of datasets used for the final deepMHCflare model.

| Dataset | Role | Observations | Positives | Pos. rate | Alleles |
| --- | --- | --- | --- | --- | --- |
| Mono- & multi-allelic training pool | Train | 167.0M | 370,879 | 0.22% | 72 |
| In-distribution validation (mono-allelic held-out) | Val (ID) | 4.0M | 6,259 | 0.16% | 23 |
| Out-of-distribution validation (21 unseen alleles) | Val (OOD) | 5.4M | 6,539 | 0.12% | 21 |
| HLA Ligand Atlas (multi-allelic patient tissues) | Val (multi) | 282.5M | 388,262 | 0.14% | 51 |

**Extended Data Table 5b:** Allele composition of each dataset split. Each row lists the alleles for one locus within one split.

| Split | Locus | *n* | Alleles |
| --- | --- | --- | --- |
| Train (72) | H-2 | 2 | D<sup>b</sup>, K<sup>b</sup> |
|  | HLA-A | 18 | 02:01, 02:03, 02:04, 02:05, 02:07, 03:01, 03:02, 03:07, 11:01, 11:10, 24:02, 24:06, 24:13, 25:01, 29:02, 29:13, 32:01, 32:02 |
|  | HLA-B | 36 | 07:02, 15:01, 15:02, 15:11, 15:15, 15:18, 18:03, 27:01, 27:02, 27:03, 27:04, 27:06, 27:07, 27:08, 27:09, 27:10, 27:16, 39:01, 39:05, 39:06, 39:09, 39:24, 41:01, 41:02, 41:03, 41:04, 41:05, 41:06, 44:05, 44:08, 44:09, 44:27, 44:28, 47:01, 51:08, 73:01 |
|  | HLA-C | 9 | 02:02, 03:03, 03:13, 05:01, 07:02, 07:04, 07:06, 12:02, 12:04 |
|  | HLA-E | 1 | 01:03 |
|  | HLA-F | 3 | 01:01, 01:03, 01:04 |
|  | HLA-G | 3 | 01:01, 01:03, 01:04 |
| Val ID (23) | HLA-A | 11 | 02:01, 02:03, 02:04, 02:05, 02:07, 03:01, 11:01, 24:02, 25:01, 29:02, 32:01 |
|  | HLA-B | 3 | 07:02, 15:01, 15:02 |
|  | HLA-C | 6 | 02:02, 03:03, 05:01, 07:02, 07:04, 12:02 |
|  | HLA-G | 3 | 01:01, 01:03, 01:04 |
| Val OOD (21) | HLA-A | 6 | 02:02, 02:06, 02:11, 11:02, 23:01, 24:07 |
|  | HLA-B | 9 | 07:04, 15:03, 15:10, 15:17, 18:01, 27:05, 44:02, 44:03, 51:01 |
|  | HLA-C | 6 | 03:02, 08:01, 08:02, 12:03, 15:02, 17:01 |
| Val multi (51) | HLA-A | 16 | 01:01, 02:01, 02:05, 03:01, 11:01, 23:01, 24:02, 25:01, 26:01, 26:08, 29:02, 30:01, 32:01, 68:01, 68:02, 69:01 |
|  | HLA-B | 21 | 07:02, 07:06, 08:01, 13:02, 14:02, 15:01, 18:01, 27:05, 35:01, 35:03, 35:08, 37:01, 39:01, 40:02, 41:01, 44:02, 44:03, 45:01, 49:01, 50:01, 58:01 |
|  | HLA-C | 14 | 02:02, 03:03, 03:04, 04:01, 05:01, 06:02, 07:01, 07:02, 07:04, 08:02, 12:03, 15:05, 16:01, 17:01 |

**Extended Data Table 6:**
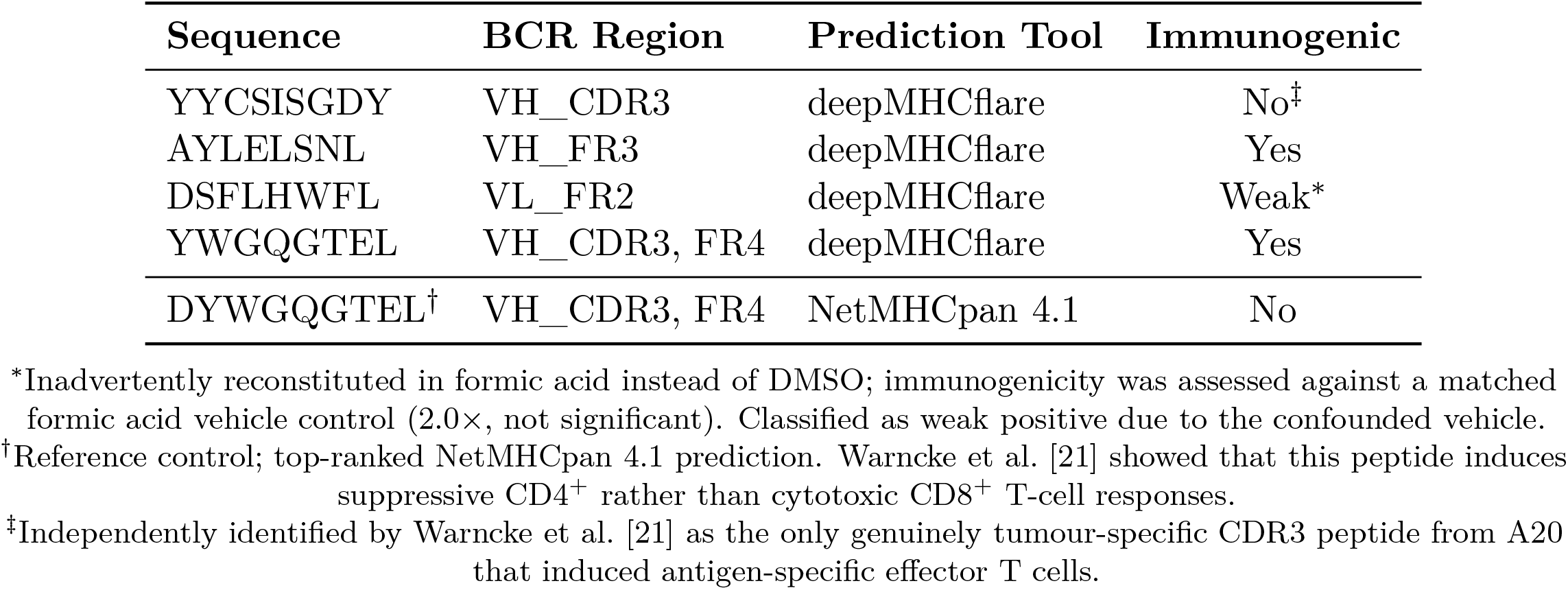
Peptide sequences tested in the prospective validation study. Four peptides were selected by deepMHCflare; DYWGQGTEL was included as a reference control (top-ranked NetMHCpan 4.1 prediction). Immunogenicity was defined as a statistically significant increase in CD8^+^ TNF-*α*^+^ T-cells (one-sided paired *t*-test, *p* < 0.05).

## Notes

### Summary of Updates

This revision corrects a submission error that affected the previous version's abstract, which did not reflect the intended text. The abstract has been updated to accurately summarize the manuscript's contributions. Beyond the abstract correction, no changes have been made to the main text, figures, methods, or supplementary materials.

