## Supplementary Note for "Resolution of recursive data corruption to transform T-cell epitope discovery"

### Supplementary Note: Out-of-distribution and multi-allelic validation

As an additional robustness check, we evaluated deepMHCflare on two held-out settings beyond the primary mono-allelic in-distribution benchmark.

**Out-of-distribution generalisation.** On the OOD validation set (21 alleles disjoint from training), deepMHCflare achieves a Precision@4 of approximately 0.5, compared to 0.8 on the in-distribution set. While performance degrades, the model retains meaningful ranking ability on alleles never seen during training, confirming that the learned representations capture transferable peptide–MHC binding patterns rather than allele-specific memorisation.

**Multi-allelic validation (HLA Ligand Atlas).** On the HLA Ligand Atlas multi-allelic dataset (245,903 contexts from 227 patient tissue samples, 51 alleles, average 5.96 MHCs per context), deepMHCflare is evaluated under clinically realistic conditions where peptide-to-allele assignment is ambiguous. The model achieves a Precision@4 of 0.28 under extreme class imbalance (all negatives sampled; positive rate $\sim$0.14%). After re-training on the full in-distribution data, Precision@4 improved to 0.36.
